# *Drosophila* ribosomal protein S5b is essential for oogenesis and interacts with distinct RNAs

**DOI:** 10.1101/600502

**Authors:** Jian Kong, Hong Han, Julie Bergalet, Louis Philip Benoit Bouvrette, Greco Hernández, Nam-Sung Moon, Hojatollah Vali, Éric Lécuyer, Paul Lasko

## Abstract

In *Drosophila melanogaster* there are two genes encoding ribosomal protein S5, *RpS5a* and *RpS5b.* Here, we demonstrate that *RpS5b* is required for oogenesis. Females lacking *RpS5b* produce ovaries with numerous developmental defects that undergo widespread apoptosis in mid-oogenesis. Females lacking germline *RpS5a* are fully fertile, but germline expression of interfering RNA targeting germline *RpS5a* in an *RpS5b* mutant background worsened the *RpS5b* phenotype and blocked oogenesis before egg chambers form. A broad spectrum of mRNAs co-purified in immunoprecipitations with RpS5a, while RpS5b-associated mRNAs were specifically enriched for GO terms related to mitochondrial electron transport and cellular metabolic processes. Consistent with this, *RpS5b* mitochondrial fractions are depleted for proteins linked to oxidative phosphorylation and mitochondrial respiration, and *RpS5b* mitochondria tended to form large clusters and had more heterogeneous morphology than those from controls. We conclude that RpS5b-containing ribosomes preferentially associate with particular mRNAs and serve an essential function in oogenesis.

## Introduction

Increasing evidence indicates that ribosomes are heterogeneous and perhaps dynamic, in contrast to the classical view of them as constitutive machinery for protein synthesis (1–11). In *Drosophila melanogaster,* nine ribosomal protein genes are each present in two paralogs, and in many of these cases one of the paralogs is primarily expressed in germline tissues (12–14). Single paralogs of four genes encoding ribosomal proteins *(RpS5b, RpS10a, RpS19b,* and *RpL22-like)* are abundantly expressed in germline stem cells and primordial germ cells (15, 16). *RpS5b* is also upregulated in *l(3)mbt* brain tumors whose cells are in an undifferentiated state and express many germline-specific genes, of which some have been implicated in tumor growth (17). These observations suggest that variant ribosomes with different protein composition may be an important factor in establishing or maintaining stem cell and/or germline identity. To investigate this, we examined the cellular and developmental functions of *RpS5b.*

## Results

### Different forms of ribosomal protein S5 are encoded by two different genes and expressed in complementary patterns

*D. melanogaster* RpS5a and RpS5b have distinct N-terminal domains of approximately 40 amino acids in length, while the remainders of the two proteins are nearly identical. The N-terminal domains of *D. melanogaster* RpS5a and RpS5b are not conserved in RpS5 orthologues in *C. elegans* or yeast, and the single mammalian form of RpS5 is a shorter protein that lacks the divergent N-terminal domain (Figure 1A). Using paralog-specific antisera that respectively recognize N-terminal peptides of RpS5a and RpS5b, we determined that both paralogs are incorporated into ribosomes, since they co-purify with a canonical ribosomal protein, RpS6 (Figure 1B). RpS5b migrates on sucrose gradients in a similar manner to RpS5a, with major peaks corresponding to the 40S small ribosomal subunit and 80S monosomes (Figure S1A). Importantly, RpS5a and RpS5b do not co-purify above background levels with each other (Figure 1B), indicating that an individual ribosome contains one isoform or the other, but not both. Both paralogs also co-purify with poly(A) binding protein (pAbp), which binds indirectly to ribosomes through interaction with translation factors, but not with a-tubulin, a negative control (Figure 1B).

**Figure 1.**
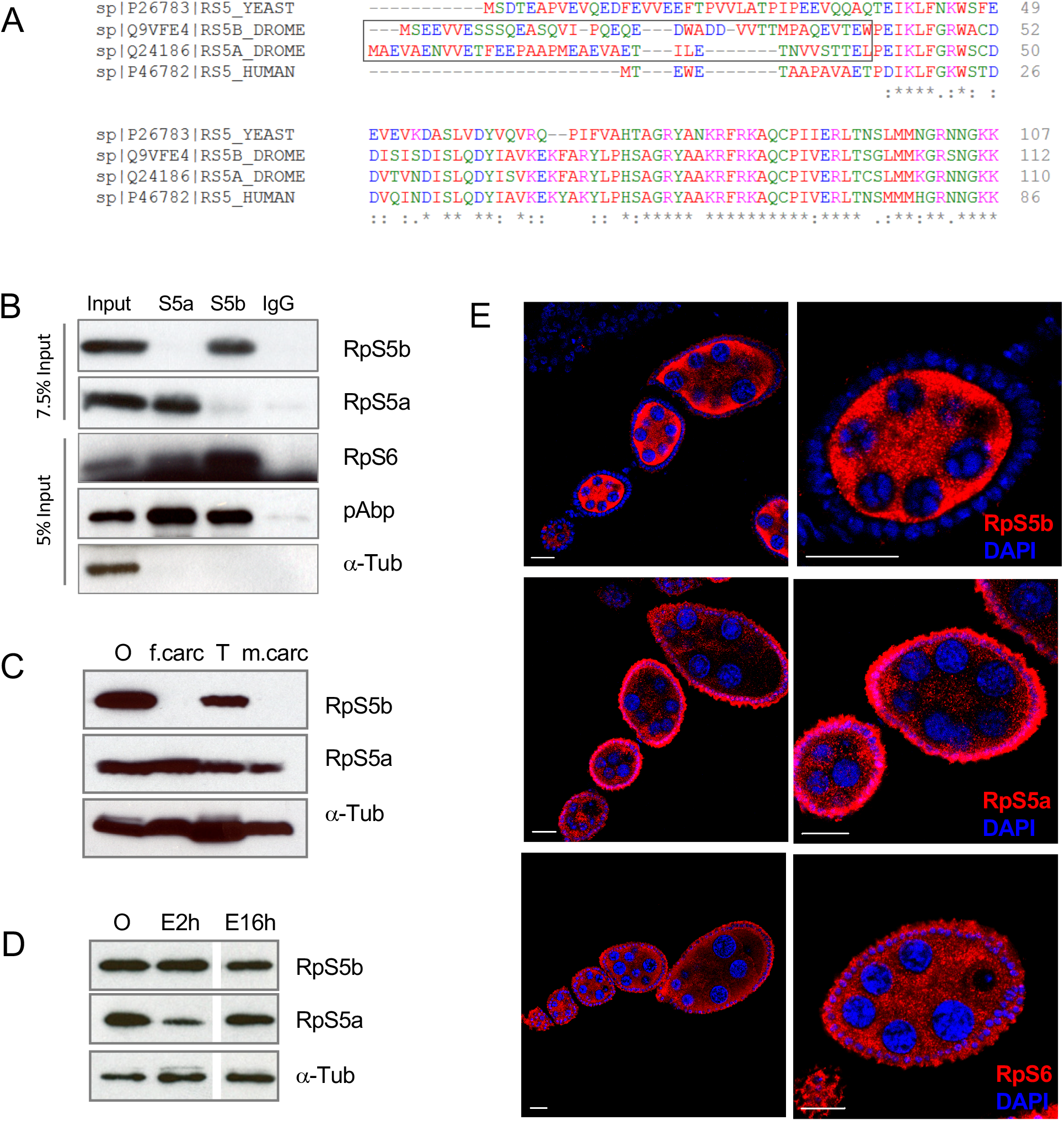
RpS5b is a germline ribosomal protein and RpS5a is expressed mostly in somatic cells. **(A)** Protein sequence alignment comparing RpS5a and RpS5b to each other and to RpS5 orthologues in yeast and human. These proteins are highly homologous but diverge in their N-terminal regions. **(B)** Co-immunoprecipitation experiments. Immunoprecipitations were conducted with antisera recognizing RpS5a, RpS5b, and non-immune IgG (top captions), blotted, and probed with antibodies as indicated on the right. **(C)** Western blot showing that RpS5b is detected in the ovary (O) and testis (T) but not in female or male carcasses (f.carc and m.carc), while RpS5a is ubiquitously expressed. **(D)** Western blot showing that RpS5b is highly abundant relative to RpS5a in 0-2h embryos (E2h), but both paralogs have relatively equal abundance in 0-16h embryos (E16h). **(E)** Immunostaining experiments showing that RpS5b is primarily expressed in the germline cells, while RpS5a is primarily, but not entirely, expressed in follicle cells. RpS6 is equally abundant in both tissues. Scale bars, 20 μm.

In adult flies, RpS5a expression is widespread, while detectable RpS5b expression is restricted to ovaries and testes (Figure 1C). Reflecting a maternal contribution of RpS5b, it is the predominant isoform in 0-2 h embryos, but in later embryos the level of RpS5a increases (Figure 1D). Immunohistochemical staining of ovaries revealed that RpS5b is mostly expressed in germline cells. RpS5a is primarily expressed in follicle cells, but some RpS5a signal above background is apparent in germline cells (Figure 1E). Similarly, in testes RpS5b is mostly expressed in germline cells while RpS5a is present in the soma (Figure S1B).

### RpS5b is specifically required for oogenesis

Mutations in canonical ribosomal protein genes including *RpS5a* (18) produce a phenotype called Minute; heterozygotes exhibit developmental delay and the adults that emerge have bristle defects and reduced viability and fertility, while homozygotes are embryonic lethal. To examine the consequences of loss of RpS5b, we obtained an *RpS5b* mutant from a large-scale insertion mutagenesis screen (*RpS5b^G5346^*) (19), subsequently referred to simply as *RpS5b,* that does not produce detectable levels of protein (Figure 2A). *RpS5b* mutant flies are fully viable and male *RpS5b* mutant flies are fertile. Female *RpS5b* mutant flies, however, are completely sterile and do not complete oogenesis. The nurse cells, follicle cells, and oocyte of these flies all differentiate, but very few egg chambers progress beyond stage 8-9, after which point apoptosis is induced, as measured by cleaved caspase-3 immunostaining (Figure 2B, C). In addition, we observed numerous developmental defects in *RpS5b* mutant ovaries. Some egg chambers contained more than 16 germ cells with one or two oocytes (Figure 2D, Figure S2A, 0.8% and 6.4% respectively in all the stage 4 and later egg chambers), some egg chambers failed to separate (Figure 2E, 8.1%), while others had 16 germ cells but two oocytes (Figure 2F, 1.1%) or a mis-localized oocyte (Figure 2G, 2.9%). We also frequently observed over-proliferation of follicle cells with multiple cell layers present at the posterior of egg chambers (Figure 2H, 54%). Polarity defects in *RpS5b* mutant oocytes were also observed. Immunostaining for a-Tubulin and for Dynein heavy chain, which marks the microtubule organizing center normally present at the posterior of the stage 7 oocyte, revealed that microtubules were improperly distributed in *RpS5b* mutant oocytes (Figure 2I-L). Consistent with this, spatial targeting of Oskar and Gurken, both microtubule-dependent processes, was disturbed in all stage 8 and later *RpS5b* oocytes that were examined (Figure 2M, N). Phalloidin staining also revealed an overabundance of F-actin in *RpS5b* oocytes (Figure S2B, C).

**Figure 2.**
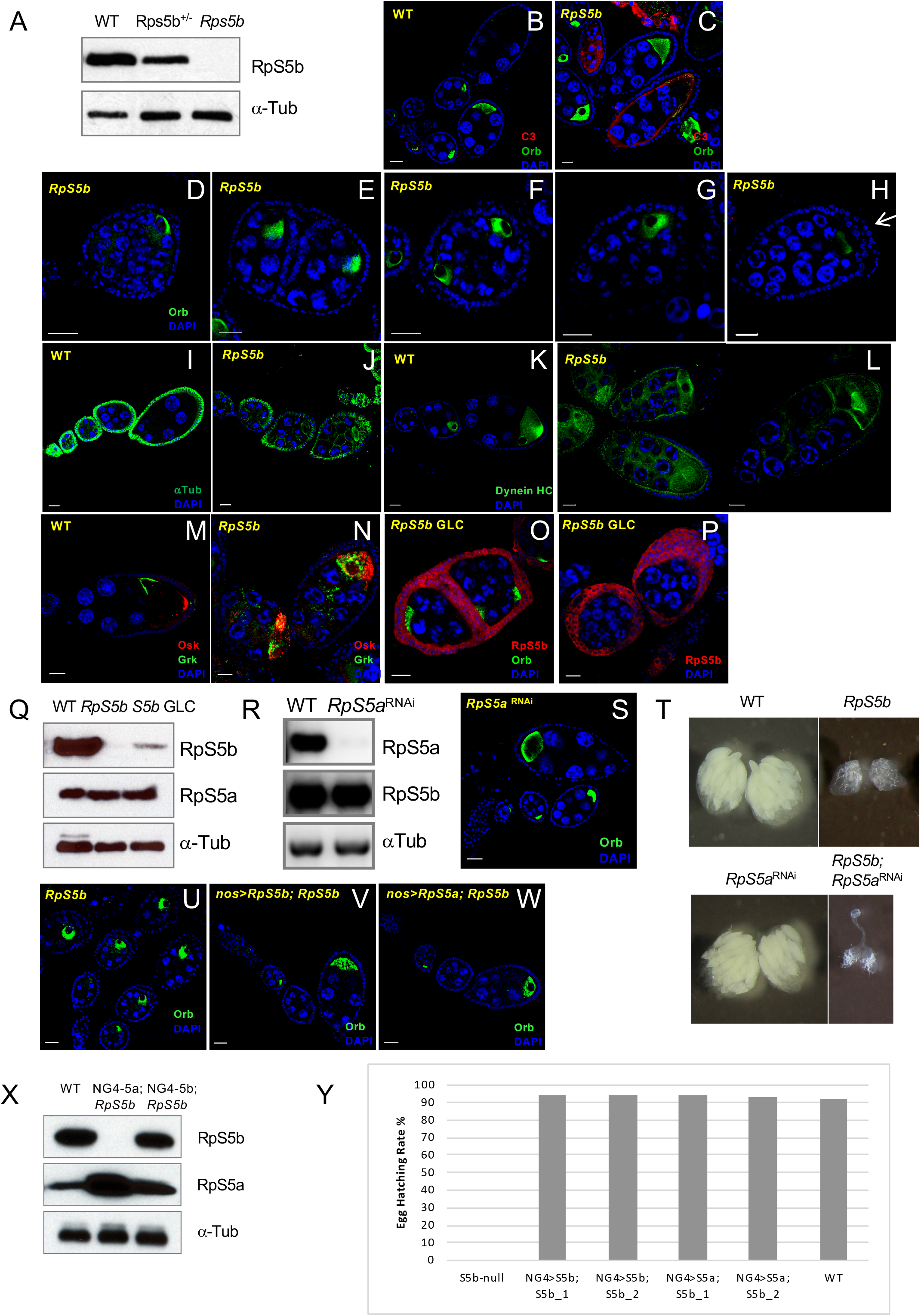
*RpS5b* mutant ovaries have numerous developmental defects that can be rescued by germline expression of *RpS5a* or *RpS5b.* **(A)** Western blot demonstrating that RpS5b is undetectable in *RpS5b* homozygotes and present at reduced levels in *RpS5b* heterozygotes, confirming the loss-of-function nature of the mutation. **(B, C)** Oogenesis does not proceed beyond stage 8 in *RpS5b* ovaries, at which point apoptosis is induced, as measured by increased levels of activated caspase-3 (C3). a-Orb is used to label the oocyte. **(D-H)** Various defects observed in *RpS5b* ovaries: **(D)** an extra round of germ cell division; **(E)** a compound egg chamber partially separated by follicle cells, **(F)** oocyte duplication in a single egg chamber; **(G)** mis-localized oocyte; **(H)** multiple layers of follicle cells at the posterior of the egg chamber. **(I-L)** Alterations in the microtubule cytoskeleton in *RpS5b* oocytes, as measured by immunostaining against **(I, J)** a-Tubulin, or **(K, L)** a-Dynein heavy chain. Note the aberrant accumulation of a-Tubulin around the oocyte in **(J)**, and the focus of Dynein in the centre of the oocyte in **(L). (M, N)** Distribution of Osk and Grk in **(M)** wildtype and **(N)** *RpS5b* oocytes, showing that deployment of these proteins is disrupted in the mutant. **(O, P)** Analysis of *RpS5b* germline clones, showing similar defects as found in the mutant, but more extreme overproliferation of follicle cells. **(Q)** Western blot showing RpS5b expression in *RpS5b* mutant and germline clones. The residual expression in the germline clones is somatic, as is also apparent in **(O, P). (R)** Western blot comparing RpS5a expression in 0-2h embryos collected from wildtype females and those expressing shRNA targeting *RpS5a* driven by the germline-specific promoter *nos*, showing the efficacy of knockdown. **(S)** Analysis of ovaries from females expressing shRNA targeting RpS5a driven by the germline-specific promoter *nos,* showing normal patterning. **(T)** Brightfield images of whole ovaries showing that *RpS5a* germline knockdown produces no phenotype on its own but worsens the *RpS5b* mutant phenotype. **(U-W)** *RpS5b* mutant ovaries **(U)** without a transgene as control or expressing transgenic **(V)** *RpS5b* or **(W)** *RpS5a* under the control of the *nos* promoter. Normal oogenesis is restored in both cases. **(X)** Western blot of lysates from 0-2h embryos collected from wildtype (WT), *nos>RpS5a; RpS5b* (NG4-5a; *S5b)* and *nos>Rps5b; RpS5b* (NG4-5b; *S5b)* females, confirming high-level expression from the transgenes. **(Y)** Graph showing hatching rates of embryos from females of the genotypes indicated, demonstrating that either *RpS5a* or *RpS5b* can fully rescue the fertility of *RpS5b* females when expressed in germline.

### Overlapping functions for RpS5a and RpS5b

Since *RpS5b* is primarily expressed in germline cells, we were surprised to observe phenotypes in the mutant that affected follicle cells as well as the germline. To analyze the requirement for *RpS5b* in the germline specifically, we used the dominant female sterile-FLP technique (20) to produce flies that lacked *RpS5b* function only in germline cells. Ovaries from *RpS5b* germline clone flies exhibited the same set of phenotypes, often with even greater severity, as the mutant flies (Figure 2O, P). In particular, many egg chambers had multiple layers of follicle cells, and encapsulation defects were observed. As measured by immunostaining (Figure 2O, P) and immunoblotting (Figure 2Q), these flies produce very little RpS5b, strictly from somatic expression, which can be visualized in follicle cells using confocal microscopy with increased gain (Figure 2O, P). Next, to investigate whether RpS5a functions in the germline, we drove expression of a short hairpin RNA that targets it in germline cells, which drastically reduced accumulation of the protein in early-stage progeny embryos (Figure 2R). Females lacking germline RpS5a were fertile and proceeded through oogenesis normally (Figure 2S, T). However, knocking down germline expression of RpS5a in an *RpS5b* mutant background gave a much more severe phenotype than the *RpS5b* mutant presents on its own; in this case flies possess only very rudimentary ovaries and no formation of egg chambers is apparent (Figure 2T). We conclude from these observations that both RpS5a and RpS5b function in germline in normal development, but also that germline RpS5b can functionally substitute for germline RpS5a.

We wanted to distinguish whether the failure of endogenous germline RpS5a to functionally substitute for RpS5b in the *RpS5b* mutant was due to functional differences between the two proteins, or whether there is insufficient endogenous expression of RpS5a in germline to rescue the function of RpS5b. To do this, we expressed both untagged and N-terminally tagged forms of RpS5a and RpS5b with *nos,* a germline specific promoter, and examined whether fertility could be restored to the *RpS5b* mutant. We observed that *nos*-driven expression of either RpS5a or RpS5b, in either untagged or Venus-tagged forms, indeed restored normal oogenesis and full fertility to the *RpS5b* mutant (Figures 2U-Y, S2D-I’). We conclude that RpS5a can substitute for RpS5b in germline tissue provided it is expressed at high enough levels.

### High-throughput analysis reveals that RpS5b associates preferentially with nuclear-encoded mRNAs involved in mitochondrial processes

As we have demonstrated, the endogenous forms of RpS5a and RpS5b are primarily expressed in different cell types, so for this reason it is very likely that they associate with different RNA populations in normal development. To investigate whether they preferentially associate with particular mRNAs when present in the same tissue, we expressed FLAG-HA-RpS5a (FH-RpS5a) or FLAG-HA-RpS5b (FH-RpS5b) in the germline using *nos*>Gal4, and then co-immunoprecipitated (co-IP) the associated RNAs with anti-FLAG antibody. We identified a set of transcripts that were enriched in FH-RpS5b co-IPs in comparison to input (Figure 3A). Gene ontology (GO) term analysis revealed that FH-RpS5b-associated mRNAs are most highly enriched for those involved in oxidative phosphorylation, electron transport chain function, electron transport chain assembly and mitochondrial translation (Figure 3B and Table S1). Far fewer mRNAs were preferentially recovered in FH-RpS5a co-IPs (Figure 3A), and no GO terms were significantly enriched among them. mRNAs enriched in FH-RpS5b co-IPs generally had very short coding regions (CDS) that were significantly different from those that associate with FH-RpS5a (Figure 3C). Taken together, these data support that there is selectivity between RpS5a and RpS5b as to the mRNAs they recruit.

**Figure 3.**
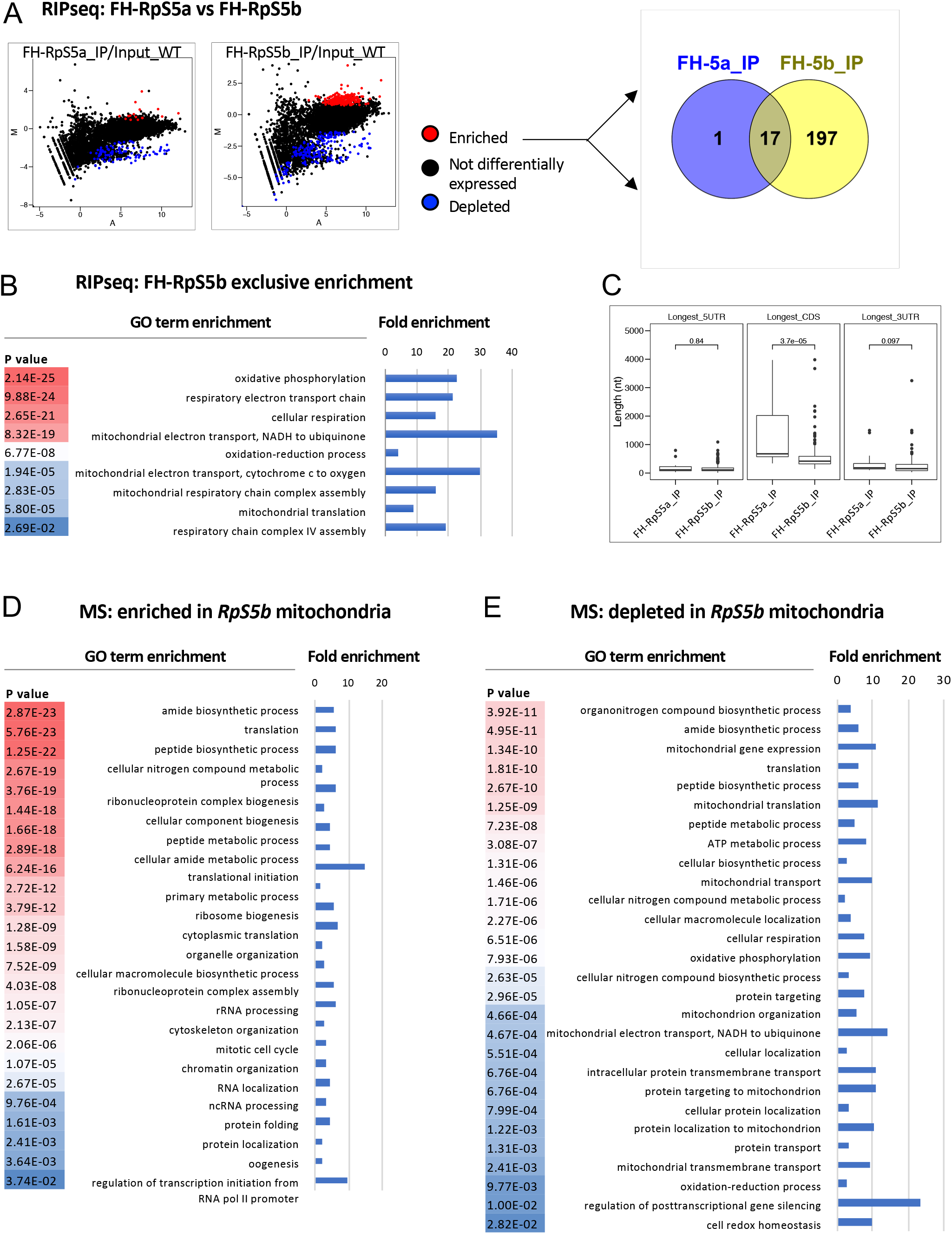
Analysis of RNA populations recruited by FLAG-HA (FH)-RpS5a and RpS5b. **(A)** MA plot of RNA immunoprecipitations using a-FLAG from ovaries with *nos*>GAL4 driven expression of either FH-RpS5a or FH-RpS5b in the wildtype background compared to input. Statistically enriched mRNAs (>2 fold, padj < 0.01) and depleted (<2 fold, padj < 0.01) are highlighted in red and blue respectively. M = *log2(pulldown)* – log2(input), A = 0.5 * *(log2(pulldown)* + log2(input)). Fold changes and adjusted p-values (padj) calculated by DESeq2 (35). The Venn diagram (http://bioinfogp.cnb.csic.es/tools/venny/) shows limited overlap between RNAs enriched in populations recruited by FH-RpS5a (FH-5a) and RpS5b (FH-5b). **(B, C)** Heat map representing biological process gene ontology (GO) terms of RNAs enriched in populations recruited by **(B)** FH-RpS5b (Statistical overrepresentation test on Pantherdb.org). The most highly significant matches are in red. The fold enrichment of each GO term is plotted in the bar chart. **(C)** Box plot of the length distribution, in nucleotides, of the 5’UTR, coding sequence (CDS) and 3’UTR for RNAs enriched in populations recruited by FH-RpS5a (FH-RpS5a_IP) and FH-RpS5b (FH-RpS5b_IP). **(D, E)** Heat maps representing the biological process GO terms associated with the proteins **(D)** enriched or **(E)** depleted in the mitochondrial fractions from *Rps5b* ovaries as compared with wild-type. The most highly significant matches are in red.

We also conducted similar co-immunoprecipitation experiments and RNA sequencing with wildtype ovaries using antisera that recognize each of the RpS5 paralogs, in order to determine whether these binding preferences are also observed in normal development. Again, largely non-overlapping sets of transcripts were enriched in either the RpS5a or RpS5b immunoprecipitates in comparison to input (Figure S3A). GO term analysis again revealed that RpS5b-associated mRNAs are most highly enriched for those involved in oxidative phosphorylation, electron transport chain function, electron transport chain assembly, and mitochondrial translation (Figure S3B and Table S2), and many RpS5b-associated mRNAs encode proteins that localize to the mitochondrial inner membrane. In contrast, only high-level GO terms related to transcriptional regulation, development, and morphogenesis were enriched among RpS5a-associated mRNAs (Figure S3B).

In addition to functional differences, we also found structural differences in the populations of RNAs that associate with RpS5a and RpS5b. RpS5b-associated mRNAs were smaller, again with significantly shorter CDS, but also with significantly shorter 5’ and 3’ UTRs, than RpS5a-associated mRNAs (Figure S3C). We also identified sequence motifs that were selectively enriched within RpS5b-or RpS5a-associated mRNA targets (Figure S3D). While these motifs have no established function, their differential enrichment provides further evidence that RpS5a and RpS5b preferentially associate with different populations of mRNAs.

To investigate whether RpS5a can be recruited to mRNAs that are normally associated with RpS5b, we examined the population of RpS5a-associated mRNAs in the *Rps5b* mutant. In this situation, the average 5’ UTR and 3’ UTR lengths of RpS5a-associated mRNAs were significantly shorter than those of RpS5a-associated mRNAs in the ovary with wildtype RpS5b, but significantly longer than those of RpS5b-associated mRNAs in the ovary with wildtype RpS5b (Figure S3C). We then individually examined the 62 mRNAs that were most enriched to RpS5b in wildtype and compared their association with RpS5a in wildtype and *RpS5b* background. We found that most of them were associated to RpS5a to an elevated degree in the *RpS5b* mutant (Figure S3E). Importantly, GO terms related to oxidative phosphorylation and mitochondrial processes were not significantly enriched among RpS5a-associated mRNAs in the *RpS5b* mutant. These results suggest that germline RpS5a can to some extent recruit mRNAs that normally associate with RpS5b when RpS5b is absent in the *RpS5b* mutant, but that this compensation is only partial and insufficient to rescue oogenesis, leading to the phenotypes we observe.

### Proteomic analysis reveals depletion of mitochondria proteins in *RpS5b* ovaries

To investigate the role of RpS5b in global translation, we used tandem mass spectrometry to compare the proteomes of similarly staged *RpS5b* and wildtype ovaries. We prepared lysates from these tissues and separated them into cytosolic and mitochondrial fractions by centrifugation. The levels of many proteins changed significantly in *RpS5b* lysates versus wildtype. We observed enrichment of some classes of proteins in the mitochondrial fraction of *RpS5b* lysates, notably those involved in translation and other RNA dependent processes (Figure 3D and Table S3). This does not necessarily imply increased levels of these proteins, as increased association of the cytosolic translational machinery with the mitochondrial outer membrane to support mitochondrial biogenesis during *Drosophila* oogenesis has been previously described (21). Consistent with the RNA analysis, proteins with GO terms related to oxidative phosphorylation were depleted in the *RpS5b* mitochondrial fraction as compared to wildtype (Figure 3E and Table S3). Proteins with higher abundance in the cytosol of *RpS5b* ovaries fell into numerous categories, while those with lower abundance included those involved in developmental processes that occur in later oogenesis, reflecting the developmental block during oogenesis in these ovaries (Figure S4).

### *Rps5b* mitochondria have altered morphology and form large aggregates

Since our analysis indicated particular effects on mitochondrial components, we examined the morphology and function of mitochondria in RpS5b-null ovaries. Immunostaining of mitochondria in the mutant ovary showed extensive clustering, in contrast to the more dispersed distribution of mitochondria in wildtype (Figure 4A). This pattern was confirmed in the *RpS5b* germline clones (Figure 4A), indicating that the mitochondrial phenotype is due to the lack of germline RpS5b.

**Figure 4.**
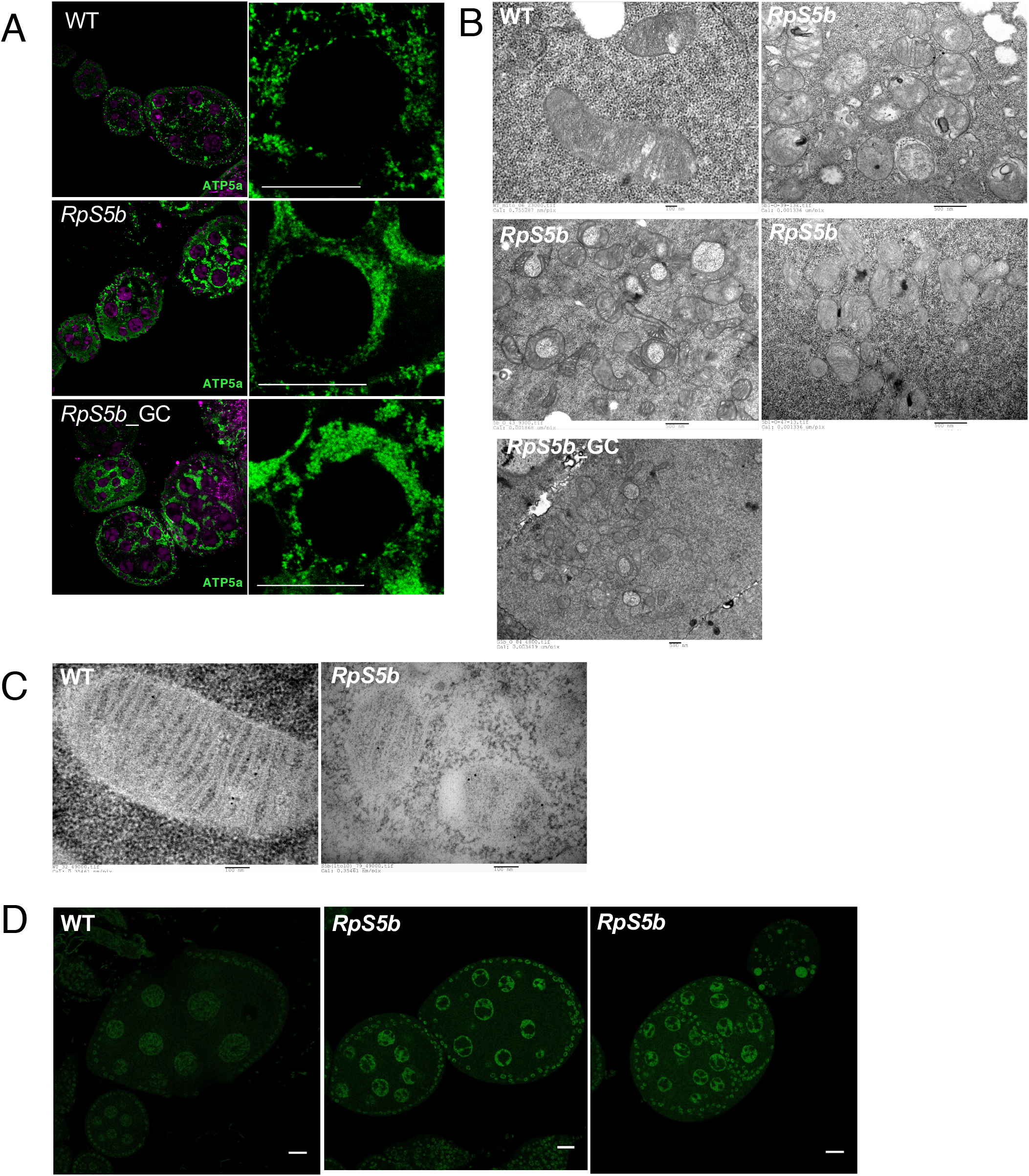
RpS5b ovaries have mitochondria with aberrant distribution and morphology, and elevated ROS. **(A)** Immunostaining for a-ATP5a, a subunit of mitochondrial ATP synthase, reveals a much more densely clustered distribution of mitochondria in nurse cells from *RpS5b,* or *RpS5b* germline clones (RpS5b_GC) than from wildtype (WT). **(B)** Transmission electron micrographs (TEM) of thin sections from nurse cells show that *RpS5b* mutant or *RpS5b* germline clones have mitochondria with aberrant cristae morphology and irregular shapes. **(C)** TEM images of mitochondria labeled with colloidal-gold conjugated a-ATP5a, illustrating morphological changes in *RpS5b* mitochondria. **(D)** Images of live ovaries from wildtype (WT) and *RpS5b* females treated with CellROX, a sensor for reactive oxygen species (ROS), showing elevated levels in *RpS5b.*

To further investigate this, we employed transmission electron microscopy (TEM) and focus ion beam scanning electron microscopy (FIB-SEM) to image the mitochondria in nurse cells of both wildtype and *RpS5b* ovaries. Again, the mitochondria in the *RpS5b* ovary appeared to form extensive clusters (Figure S5A). In most cases, *RpS5b* mitochondria were less elongated and had less well resolved cristae (Figure 4B, C). Sometimes two *RpS5b* mitochondria adhere to each other via the outer membrane (Figure 4B), and some *RpS5b* mitochondria bend and engulf some cytoplasm, forming a donut shape. These patterns were observed both in *RpS5b* mutant and germline clones (Figure 4B). To further analyze the structure of *RpS5b* mitochondria, focused ion beam-scanning electron microscopy (FIB-SEM) of 120 consecutive serial sections of 4 nm thickness was performed. Reconstructed data produced in this way revealed the 3D structure of the donut-shape mitochondria as engulfing cytosolic material (Figure S5B, movie showing z-stack). To test whether loss of RpS5b affects mitochondrial function in vivo, we treated live ovaries with CellROX, a reagent that measures reactive oxygen species (ROS) levels. We observed markedly elevated ROS levels in *RpS5b* mutant ovaries, consistent with mitochondrial dysfunction (Figure 4D).

## Discussion

Our experiments demonstrate that RpS5b, one of two RpS5 paralogs in *Drosophila,* is required for completion of oogenesis and for female fertility. While RpS5a is primarily expressed in somatic cells and RpS5b is primarily expressed in germline, we present genetic evidence that both paralogs function in germline in early stages of oogenesis, as germline-specific knockdown of *RpS5a* exacerbates the *RpS5b* phenotype, producing a very early arrest in oogenesis. There is other evidence that non-canonical components of the translational machinery is required in pluripotent cells. For example, two forms of eukaryotic initiation factor 4G (eIF4G) are expressed in *Drosophila* testes. In a situation analogous to RpS5a and RpS5b, eIF4G is primarily expressed in soma and eIF4G2 is primarily expressed in germline, but germline specific knockdown of both eIF4G and eIF4G2 produces a more severe phenotype than germline specific knockdown of eIF4G2 alone (22). Another example is in *Drosophila* oogenesis, where Mextli, a protein related to eIF4G, is expressed specifically in germline stem cells and early-stage cystocytes, where it binds eIF4E and promotes translation (23). It is also noteworthy that, while most mammalian ribosomal proteins exist only in one form, an exception is RpS4, which is encoded by three genes, one on the X and two on the Y chromosome, with one of the Y-linked genes primarily expressed during spermatogenesis (24). Taken together, these results indicate that germline cells rely upon a specialized translational machinery.

We further demonstrate that ribosomes containing RpS5a and RpS5b associate with distinct populations of RNAs. Most notably, RpS5b-associated mRNAs are highly enriched for those encoding mitochondrial proteins involved in oxidative phosphorylation. Consistently, this class of proteins is depleted in *RpS5b* ovaries, which also exhibit abnormalities in mitochondrial distribution and morphology. Rapid growth in later stages of oogenesis requires adequate energy production and a high level of mitochondrial biogenesis (21), so the mitochondrial abnormalities we observe in *RpS5b* ovaries may explain some aspects of their phenotype. Mitochondrial dysfunction has been previously linked by others to female infertility and failure of oogenesis, including in *Drosophila* (25).

In conclusion, our study provides further direct evidence that specialized ribosomes can play a pivotal physiological role in a metazoan. In yeast where many ribosomal proteins are encoded by paralog pairs, three specific paralogs (Rpl1b, Rpl2b, and Rps26b) are required for proper mitochondrial morphology and function (26). We therefore conclude that translational control through variant ribosomes is a well conserved regulatory mechanism that is particularly important for ensuring appropriate expression of nuclear genes encoding mitochondrial proteins.

## Materials and Methods

### Immunostaining

Immunostainings were carried out as described (27) with the following modifications. Ovaries were dissected from 5-10 day old female flies in PBST (PBS with 0.3% Triton X-100) and fixed in PBST with 4% formaldehyde for 20 min at room temperature (RT). Fixed ovaries were washed at least 3 times with PBST, permeabilized in PBS with 1% Triton X-100 for 1 h, and blocked in PBSTA (PBST with 1% bovine serum albumin) for 1 h. Samples were incubated with primary antibodies overnight at 4°C in PBSTA. The dilutions of the primary antibodies were as follows: a-RpS5b (peptide antibody generated by Biomatik) 1:1000; a-RpS5a (peptide antibody was generated by Biomatik) 1: 1000); a-aTubulin (Sigma) 1:5000; a-Orb (DSHB) 1:50; a-Dhc (DSHB) 1:50; a-cleaved Caspase 3 (Abcam), 1:200; a-ATP5A (Abcam) 1:1000; a-Aub 1:1000; a-Osk 1:500; a-Grk 1:500. Antisera were produced in the Lasko lab unless otherwise noted. Venus-tagged proteins were imaged directly under ultraviolet light.

Samples were washed and incubated in the dark with fluorescent secondary antibody (preadsorbed goat anti-rabbit Alexa Fluor555, or goat anti-mouse Alexa Fluor488, Molecular Probes, 1:500) in PBSTA for 90 min at RT. Samples were then dark washed and counterstained with DAPI, mounted in 1% DABCO (in 90% glycerol) anti-fade reagent, and examined under confocal microscopy (Zeiss LSM510).

### Immunoprecipitation and Western blot analysis

20 μl Dynabeads (Life Technologies) was conjugated to 2 μg primary antibody in 100 μl of PBS with 1% NP-40 for 30 min at RT. Ovaries were dissected in PBST, immediately transferred to ice and lysed in 50 mM HEPES, pH 7.5, 100 mM KCl, 12 mM MgCl_2_, 1% NP-40, 1mM dithiothreitol, 1x Halt protease inhibitor, RNaseOut RNase inhibitor and 100 μg/mL cycloheximide (lysis/IP buffer) on ice. The lysate was centrifuged at 10,000 x g for 15 min at 4°C. Supernatant was transferred to the Ab-conjugated beads, and incubated on a rotator at 4°C for 1.5 h. The beads were washed with lysis/IP buffer and eluted with 2x SDS loading buffer by boiling for 3 min at 95°C. Protein samples were resolved on SDS-PAGE and probed with antibodies. Primary antibody concentrations used in Western blots were: α-RpS5a 1:1000; α-RpS5b 1:1000; α-RpS6 (Cell Signalling) 1:300; α-αTubulin 1:5000; α-pAbp 1:30,000; α-ND-30 1:1000; α-Porin (Abcam) 1:500.

### Sucrose gradients

Lysates were prepared from wild-type ovaries and fractionated on 10-50% linear sucrose gradients. Fractions were run on an SDS-PAGE gel, immunoblotted, and incubated with antisera recognizing proteins as indicated. a-Tubulin was used as a control cytosolic protein.

### Cell lysate fractionation and mitochondria purification

Mitochondria were purified as described (28) with the following adaptations. Dissected ovaries were homogenized in mitochondria isolation buffer (250 mM sucrose, 10 mM Tris, pH 7.5, 1 mM EDTA, 1x Halt protease inhibitor). The crude lysate was centrifuged at 600 x g for 7 min at 4°C. The pellet was discarded and the supernatant was centrifuged at 10,000 x g for 15 min at 4°C. The supernatant was the cytoplasmic S10 fraction. The pellet containing mitochondria was washed and resuspended in 10 mM HEPES and solubilized by sonication.

### RIPseq

Polysome immunoprecipitations were carried out as described (29) with the following adaptations. 15 mg of ovaries were homogenized using a Dounce in 10% w/v polysome buffer (50 mM Tris, pH 7.5, 100 mM KCl, 12 mM MgCl_2_, 1% NP-40, 1 mM DTT, 1 mg/mL heparin, 200 U/mL RNaseOut RNase inhibitor, protease inhibitors and 100 μg/mL cycloheximide) and centrifuged at 10,000 rpm for 10 min. 50 μl protein A/G magnetic beads (Dynabeads, Invitrogen) were saturated with 5 μg BSA and 200 U/mL RNaseOut, and homogenates were pre-cleared on 20 μl of beads for 1 h at 4°C. Lysates, beads and 5 μg of a-RpS5b, a-RpS5a or non-immune IgG antibodies were mixed and rotated for 3 h at 4°C. The beads were then washed 5 times in high salt buffer (50 mM Tris pH 7.5, 300 mM KCl, 12 mM MgCl_2_, 1% NP-40, 1 mM DTT, 100 μg/mL cycloheximide). 25% of the washed beads were saved for Western blot. The bound RNA was purified by addition of 2.5 volumes of RLT buffer (Qiagen) and extracted using an RNeasy Mini kit according to the manufacturer’s instructions (Qiagen). RNA concentrations were measured with a Nanodrop (ThermoFisher) and quality was evaluated with a Bioanalyser (Agilent).

### In silico analysis of RNA sequencing

Read quality was assessed using FastQC. Read alignment was executed using TopHat on the *Drosophila* BDGP5.78/dm3 genomes from Ensembl (30). Read count was obtained with featureCounts (31). Normalized count values and differential expression was computed with DESeq2 (32). Longest isoform UTR and CDS lengths were obtained through the R biomaRt library (33). Motif enrichment was determined using HOMER (34).

### Proteomics

Purified mitochondria and the cytoplasmic S10 were subjected to liquid chromatography–tandem MS (LC–MS/MS) (Proteomics core facility, IRIC). See figure legends for the details of data analysis.

### TEM Imaging

Ovaries were dissected from 5-15 day old female flies, fixed in 2.5% (vol/vol) glutaraldehyde in 0.1 M sodium cacodylate buffer, pH 7.4, and incubated overnight at 4°C. The samples were washed with sodium cacodylate buffer three times for 30 min, and then post-fixed in 0.1 M sodium cacodylate buffer containing 1% (wt/vol) OsO4 and 1.5% (wt/vol) potassium ferrocyanide for 2 h at 4°C and en bloc stained with 1% (wt/vol) aqueous uranyl acetate at 4 °C. Samples were dehydrated in five successive steps of acetone and water [30–90% (vol/vol)], each for 15 min at room temperature followed by 100% acetone (3× 20 min). The samples were incubated with increasing concentrations [30–100% (vol/vol)] of low viscosity EPON 812 replacement (Mecalab Limited, Montreal, QC) and acetone over a period of 24 h, and then polymerized at 65 °C for 48 h. Ultrathin sections (70-100 nm) were cut from the resin blocks using a Leica Microsystems EM UC7 ultramicrotome (Leica Microsystems) with a Diatome diamond knife (Diatome Ltd, Nidau, Switzerland). The sections were transferred onto 200-mesh Cu TEM grids (EMS, Hatfield, PA) and poststained with 4% (wt/vol) aqueous uranyl acetate for 8 min followed with Reynold’s lead for 5 min. Sections were imaged with an FEI Tecnai 12 TEM (Thermo Fisher Scientific, Hillsboro, OR) equipped with an AMT XR80C CCD camera (Advanced Microscopy Techniques, Woburn, MA) at an accelerating voltage of 120 kV in bright-field mode.

### Serial Block Face Imaging

Sample blocks for 3D characterization by FIB-SEM were prepared as described above for TEM. The blocks were trimmed with a razor blade to expose the region of interest (ROI), mounted on fixed 45° pretilt SEM stubs and coated with a 2 nm layer of platinum using a Leica Microsystems EM ACE600 sputter coater (Leica Microsystems) to enhance electrical conductivity. Milling of serial sections and imaging of the block face after each z-slice was carried out with the Helios Nanolab 660 DualBeam using Auto Slice & View G3 ver 1.2 software (Thermo Fisher Scientific). The sample block was first imaged to determine the orientation of the block face and ion and electron beams. A 2 μm layer of platinum was deposited on the surface of the ROI to protect the resin volume from ion beam damage and to correct for stage and/or specimen drift, i.e. orthogonal to the block face of the volume to be milled. Trenches on both sides of the ROI were created to minimize re-deposition during automated milling and imaging. Fiducials were generated for both ion and electron beam imaging and used to dynamically correct for drift in the x- and y-directions during data collection by applying appropriate SEM beam shifts. Milling was carried out at 30 kV with an ion beam current of 2.5 nA, stage tilt of 4°, and working distance of 4 mm.

At each step, a 4-nm slice of the block face was removed by the ion beam. Each newly milled block face was imaged with the in-column detector (ICD) at an accelerating voltage of 2 kV, beam current of 0. 4 nA, stage tilt of 42°, and working distance of 2.5 mm. The pixel resolution was 3.9 nm with a dwell time of 30 μs per pixel. Pixel dimensions of the recorded image were 3072 x 2048 pixels. Three hundred fifty images were collected and the image contrast inversed. Visualization and direct 3-D volume rendering of the acquired datasets was performed with Amira for Life Sciences software (Thermo Fisher Scientific) with 100 successive images selected based on the ROI, i.e., mitochondria.

## Acknowledgements

We are grateful to Hana Antonicka and Eric Shoubridge for helpful discussions and assistance with mitochondrial fractionations. This work was supported by CIHR grants MOP-44050 and IOP-107945 to P. L. J. B. was supported by a FRQ-S postdoctoral scholarship, E. L. is a Junior 2 FRQ-S Scholar. Work in the Lécuyer lab is supported by an FRQ-S strategic team grant and by CIHR grant MOP-137096.

## Supplemental Figure Legends

**Figure S1.**
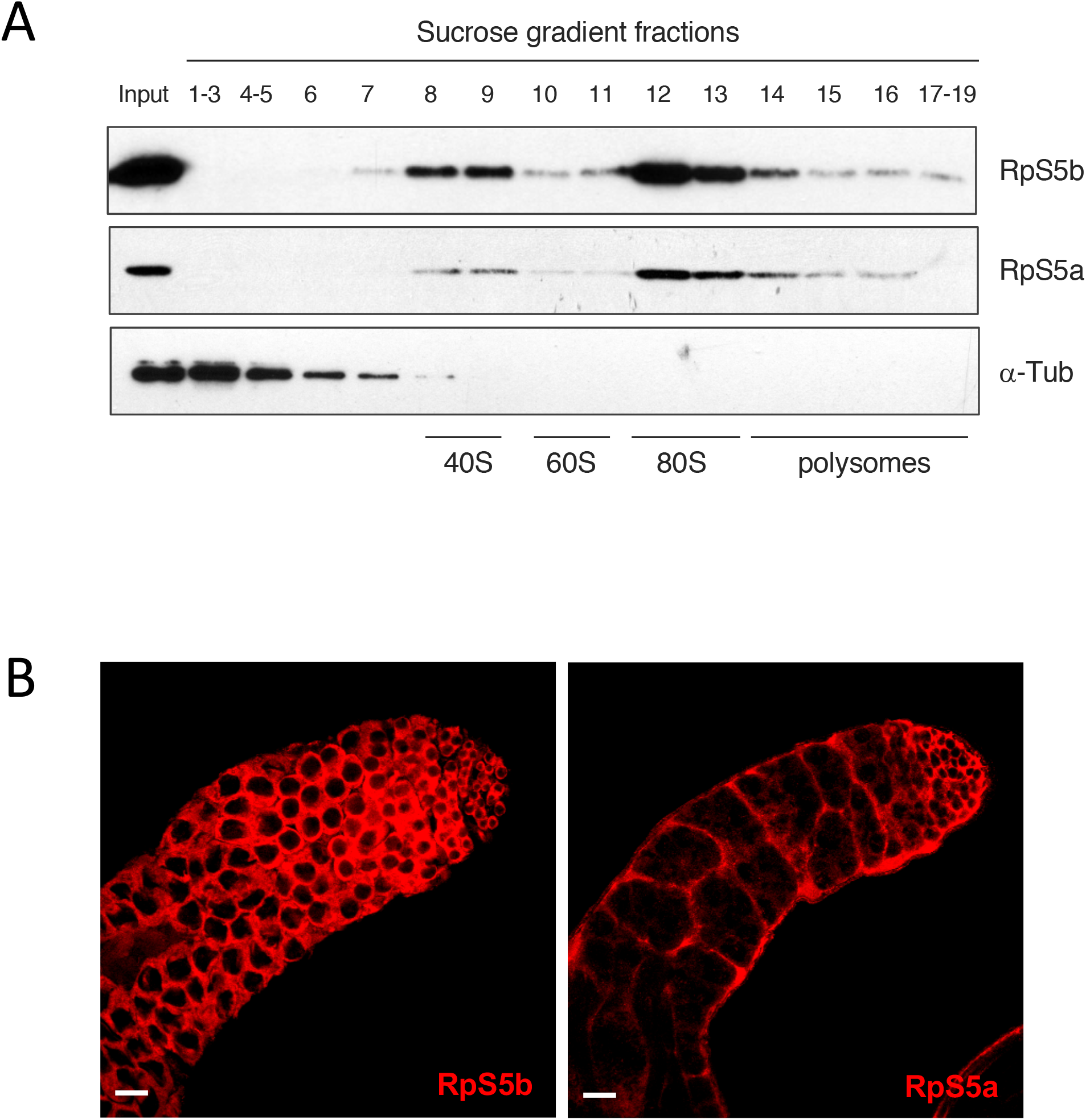
**(A)** RpS5a and RpS5b cosediment with canonical ribosomal protein RpS6 in sucrose gradients. Lysates were prepared from wildtype embryos and fractionated on 15-50% sucrose gradients. Fractions were run on an SDS-PAGE gel, immunoblotted, and incubated with antisera recognizing proteins as indicated. a-Tubulin was used as a control cytosolic protein. **(B)** RpS5a and RpS5b have complementary expression patterns in testes. Immunostaining of whole-mount testes indicates that RpS5a is primarily expressed in somatic cells while RpS5b is primarily expressed in germline.

**Figure S2.**
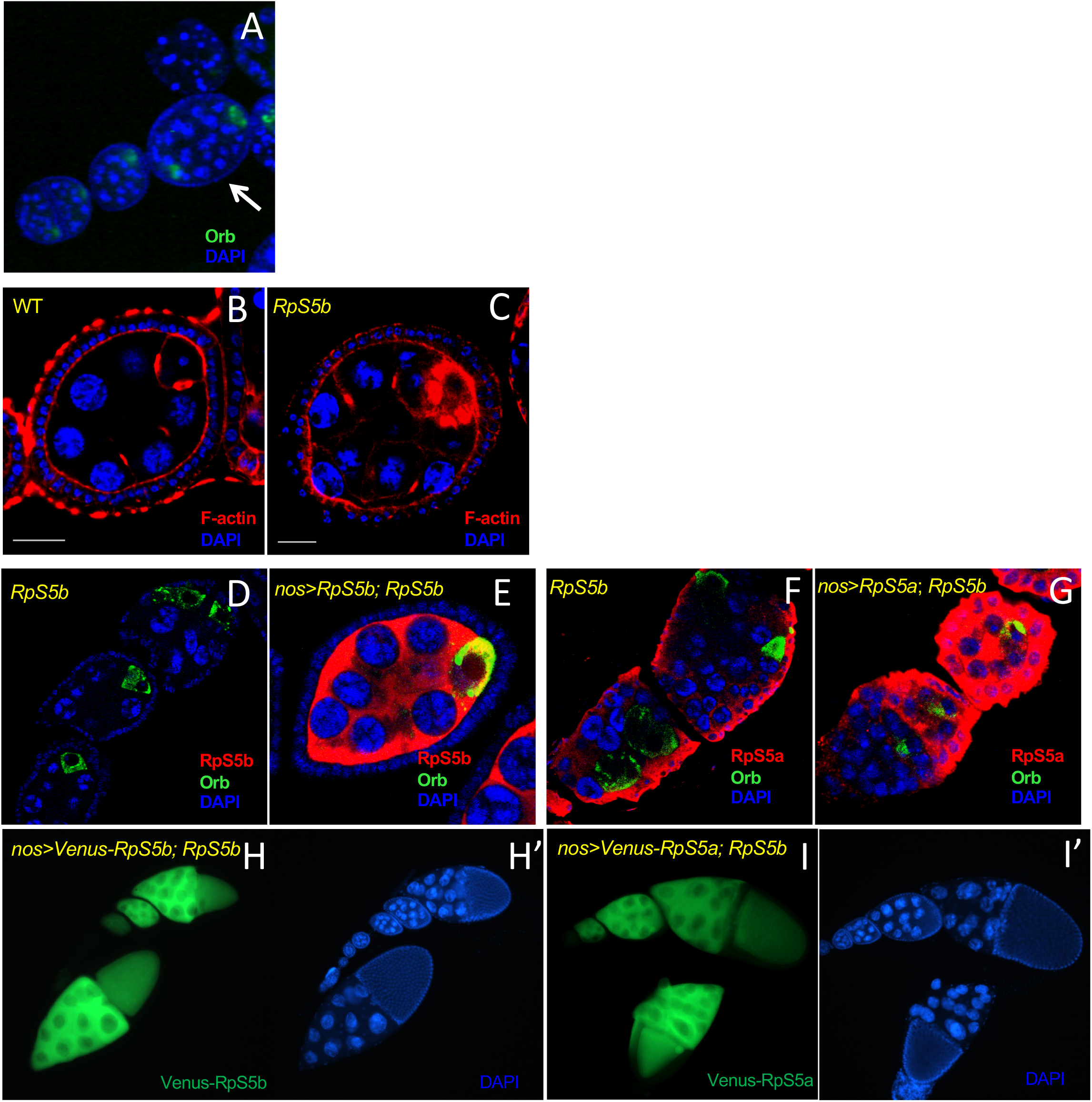
**(A)** DAPI and Orb staining of *RpS5b* ovaries shows an egg chamber with supernumerary nurse cells and two oocytes. **(B, C)** Rhodamine-conjugated phalloidin staining of **(B)** wild-type and **(C)** *RpS5b* ovaries shows excessive accumulation of F-actin in *RpS5b* oocytes. **(D-I’)** Transgenic expression of either RpS5 isoform in germline rescues normal morphology. **(D, E)** *nos*-driven expression of untagged RpS5b in germline in the *RpS5b* mutant rescues normal morphology and posterior localization of the oocyte, marked with Orb. **(F, G)** *nos*-driven expression of untagged RpS5a in germline in the *RpS5b* mutant rescues normal morphology and posterior localization of the oocyte, marked with Orb. **(H, H’)** *nos*-driven expression of Venus-tagged RpS5b in the *RpS5b* mutant rescues late stages of oogenesis. **(I, I’)** *nos*-driven expression of Venus-tagged RpS5a in the *RpS5b* mutant rescues late stages of oogenesis.

**Figure S3.**
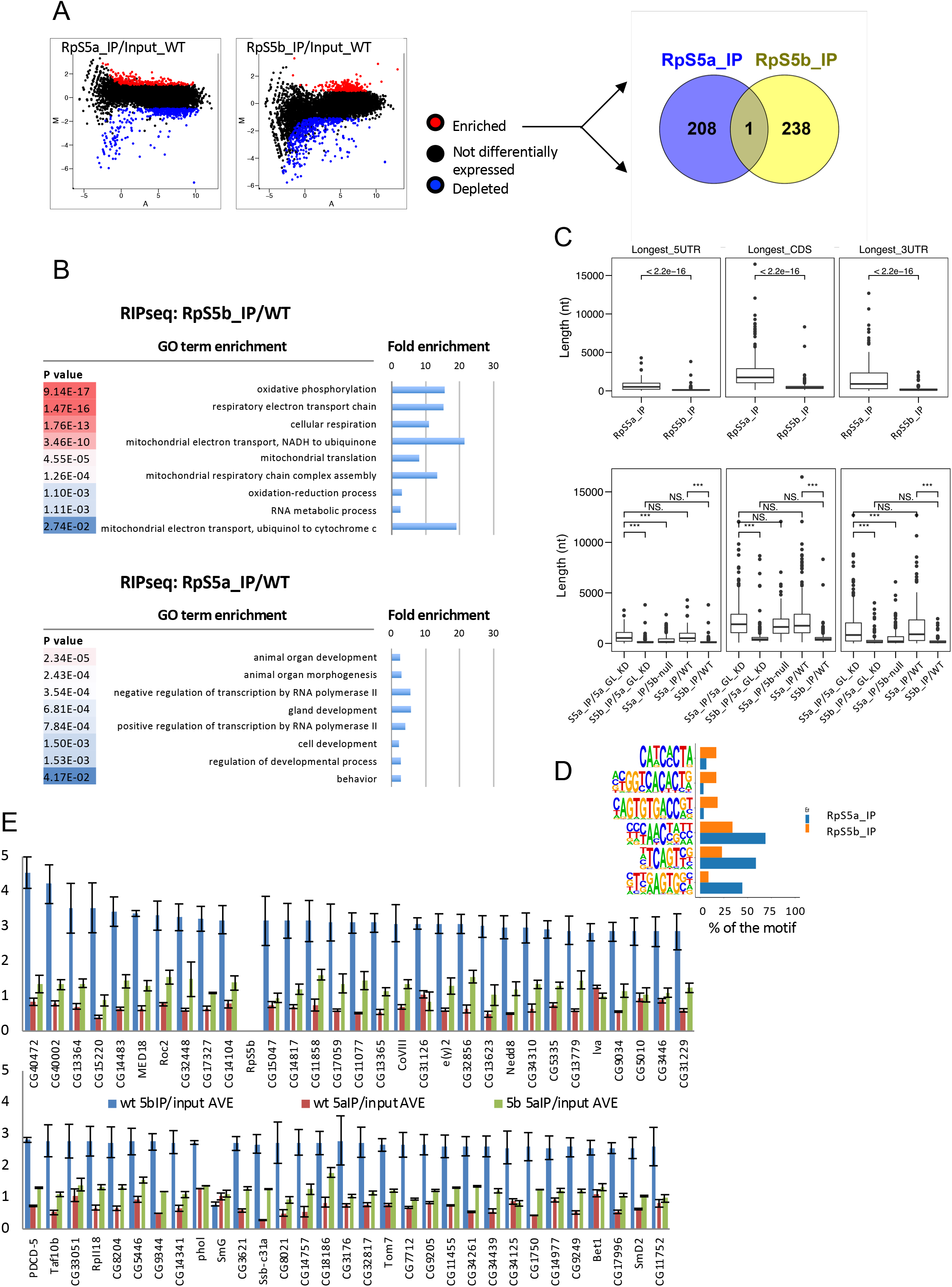
**(A)** MA plot of RNA immunoprecipitations from wildtype ovaries with a-RpS5a (RpS5a_IP) or a-RpS5b (RpS5b_IP) compared to input. Statistically enriched (>2 fold, padj < 0.01) and depleted (<2 fold, padj < 0.01) are highlighted in red and blue respectively. M = *log2(pulldown)* – log2(input), A = 0.5 * *(log2(pulldown)* + log2(input)). Fold changes and adjusted p-values (padj) calculated by DESeq2 (35). The Venn diagram (http://bioinfogp.cnb.csic.es/tools/venny/) shows minimal overlap between RNAs enriched in populations recruited by RpS5a and RpS5b. **(B)** Heat map representing biological process gene ontology (GO) terms of RNAs enriched in populations recruited by RpS5b and RpS5a, respectively. The most highly significant matches are in red. The fold enrichment of each GO term is plotted in the bar charts. **(C)** Box plot of the length distribution, in nucleotides, of the 5’UTR, coding sequence (CDS) and 3’UTR for RNAs enriched in populations recruited by RpS5a (RpS5a_IP) and RpS5b (RpS5b_IP) (upper panel). Box plots showing the sizes of 5’ UTRs, coding sequences (CDS), and 3’ UTRs of RNAs associated with RpS5a when RpS5a is expressed under *nos* control (S5a_IP/5a_GL_KD), RNAs associated with RpS5b when RpS5a is expressed under *nos* control (S5b_IP/5a_GL_KD), RNAs associated with RpS5a in the *RpS5b* mutant (S5a_IP/5b-null), as well as RNAs associated with RpS5a and RpS5b in the WT (S5a_IP/WT and S5b_IP/WT, respectively) (lower panel). **(D)** Enriched motifs in RpS5a-associated RNAs (blue) and RpS5b-associated RNAs (orange) as computed by HOMER (34). **(E)** RpS5b-associated RNAs are recruited to RpS5a in the absence of RpS5b. Comparison of mRNA abundance in co-immunoprecipations from wildtype ovaries with RpS5b (blue), RpS5a (red), or with RpS5a in the *RpS5b* mutant (green). Virtually all these RNAs are more abundant in RpS5a immunoprecipitations from the *RpS5b* mutant than from wildtype.

**Figure S4.**
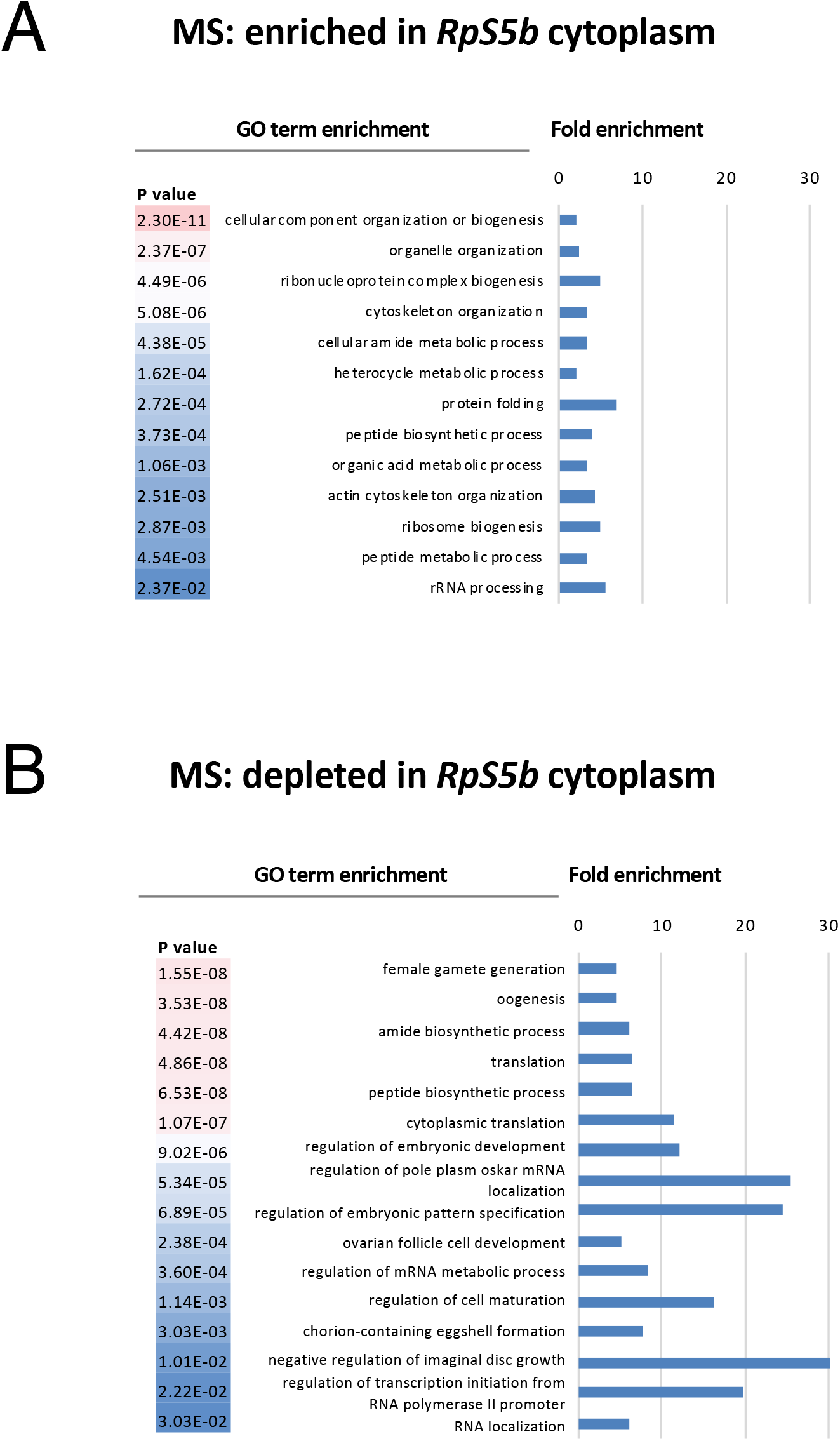
Heat maps representing the biological process GO terms associated with the proteins **(A)** enriched or **(B)** depleted in the cytosolic fractions from *Rps5b* ovaries as compared with wild-type. The most highly significant matches are in red. The fold enrichment of each GO term is plotted in the bar charts.

**Figure S5.**
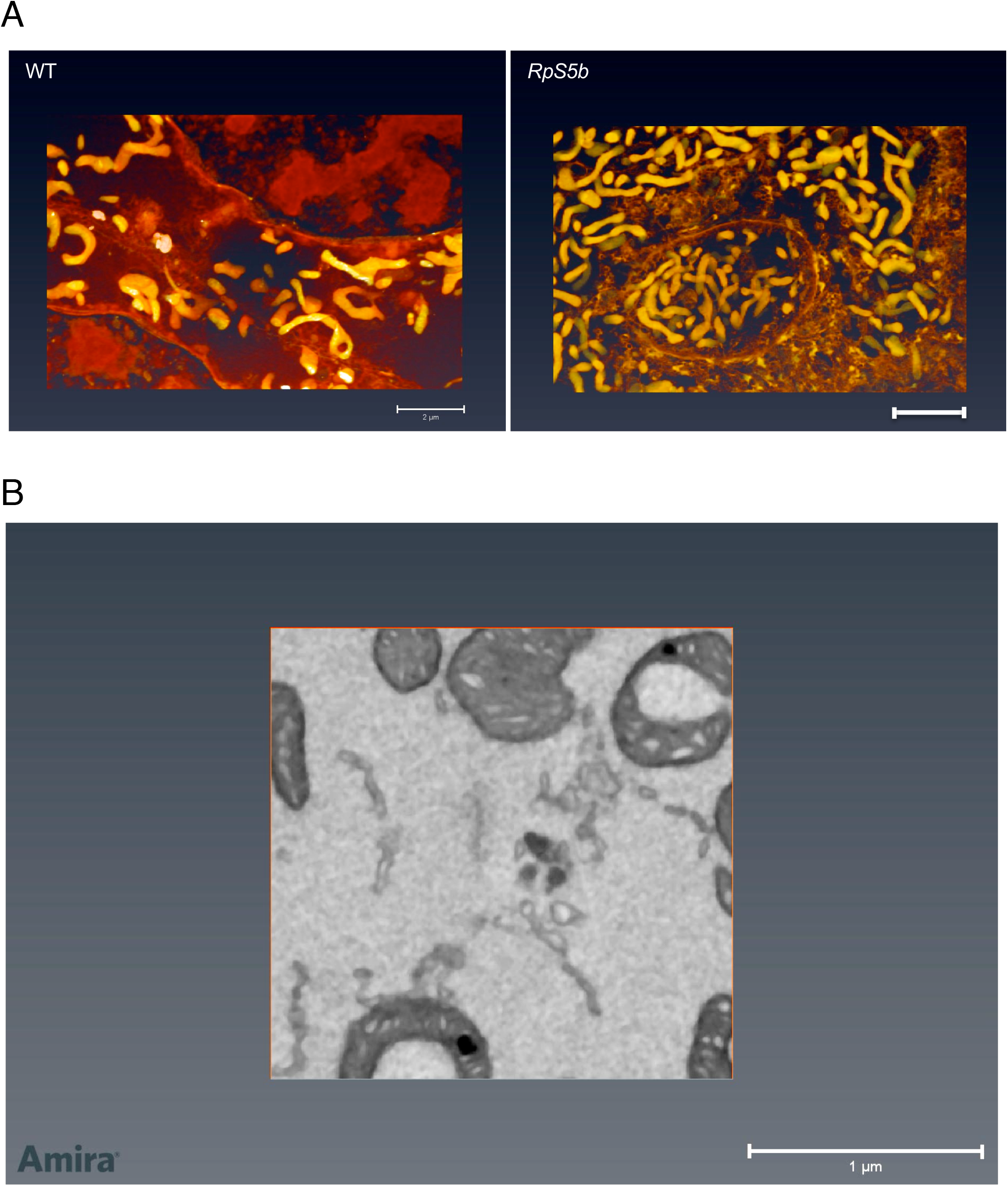
*RpS5b* mitochondria cluster more densely and have altered morphology. **(A)** FIB-SEM reconstructions showing mitochondrial distributions in wildtype (WT) and *RpS5b* nurse cells. **(B)** Movie showing z-stack reconstruction illustrating a donut-shaped mitochondrion engulfing other cytosolic material in an *RpS5b* nurse cell.

### Supplemental Tables

**Table S1**. Complete lists of mRNAs enriched in a-FLAG immunoprecipitations of *nos*>GAL4 FH-RpS5a and *nos*>GAL4 FH-RpS5b, along with GO term analysis for overrepresented biological processes carried out using PANTHER (36). Relevant to Figure 3A-C.

**Table S2**. Complete lists of mRNAs enriched in immunoprecipitations of endogenous RpS5a and RpS5b, along with GO term analysis for overrepresented biological processes carried out using PANTHER (36). Relevant to Figure S3.

**Table S3**. Complete lists of proteins enriched or depleted in mitochondrial and cytosolic fractions from *RpS5b* ovaries as compared with wild-type ovaries, along with GO term analysis for overrepresented biological processes carried out using PANTHER (36). Relevant to Figure 3D-E and Figure S4.

